# A subgenome-resolved and chromosome-scale reference genome assembly of allotetraploid wheat wild relative *Aegilops peregrina*

**DOI:** 10.64898/2026.08.28.747929

**Authors:** Jatinder Singh, Santosh Gudi, Peter J. Maughan, Upinder Gill, Rajeev Gupta

## Abstract

*Aegilops peregrina* is a wild allotetraploid wheat wild relative and an important source of genetic diversity for stress tolerance and agronomic traits. Here, we report a subgenome-resolved, chromosome-scale reference genome assembly of a drought tolerant and stem rust resistant *Ae. peregrina* accession PI 604178 generated using PacBio HiFi and Hi-C sequencing. The 10.13 Gb assembly contains 98.81% of sequence anchored to 14 pseudomolecules representing the seven Sᵖ and seven Uᵖ chromosomes, with contig and scaffold N50 values of 25.84 and 746.48 Mb, respectively. The assembly achieved a consensus quality value of 74.61, 97.83% k-mers completeness, and 99.9% BUSCO completeness. LTR Assembly Index values of 20.43 and 18.79 for the Sᵖ and Uᵖ subgenomes, respectively, further supported high continuity across repeat-rich regions. Repetitive elements comprise 85.93% of chromosome-anchored assembly. We annotated 59,910 high-confidence protein-coding genes, with comparable gene representation across the two subgenomes. This reference genome provides a high-quality genomic framework for comparative analyses, characterization of important loci regulating agronomic and resilience related traits, and sequence-guided exploitation of *Ae. peregrina* allelic diversity for wheat improvement.

## Background & Summary

Bread wheat (*Triticum aestivum* L.) provides approximately one-fifth of the calories consumed globally^1^. But its genetic diversity is constrained by the limited number of hybridization events involved in the origin of hexaploid wheat^2–4^. Wheat wild relatives in the genus *Aegilops* represent an important reservoir of genetic diversity for wheat improvement^5–7^. The genus comprises 23 species encompassing diploid C, D, M, N, S, and U genome lineages and several polyploid combinations^8,9^. *Aegilops peregrina* (Hack.) Maire & Weiller (syn. *Ae. variabilis* Eig) is an allotetraploid (2n=4x=28) with genome constitution S^p^S^p^U^p^U^p^, combining a U genome from *Ae. umbellulata* with S genome of Sitopsis origin^10^. This species is an important tertiary gene pool donor for wheat improvement. Genetic variation from *Ae. peregrina* has been used to introduce resistance to leaf rust^11,12^, stripe rust^13,14^, powdery mildew^15^, cereal cyst nematode^16^, and root knot nematode^17^. Notably examples included, leaf rust resistance genes *Lr59*, *LrP*, and *LrAp*, stripe rust resistance gene *YrP* and nematode resistance genes *CreX*, *CreY*, and *Rkn*^9^. *Ae. peregrina* has also been found to be tolerant to abiotic stresses such as drought^18^ and salinity^19^ and it has been reported to have increased grain protein and micronutrient content^9^. This species contains substantial resistance to highly virulent stem rust races^20^. *Ae. peregrina* accession PI 604178 was selected in this present study based on its seedling drought tolerance response (Fig. 1A) and resistance to stem rust races^20^ including TTKSK and TRTTF.

**Fig. 1:**
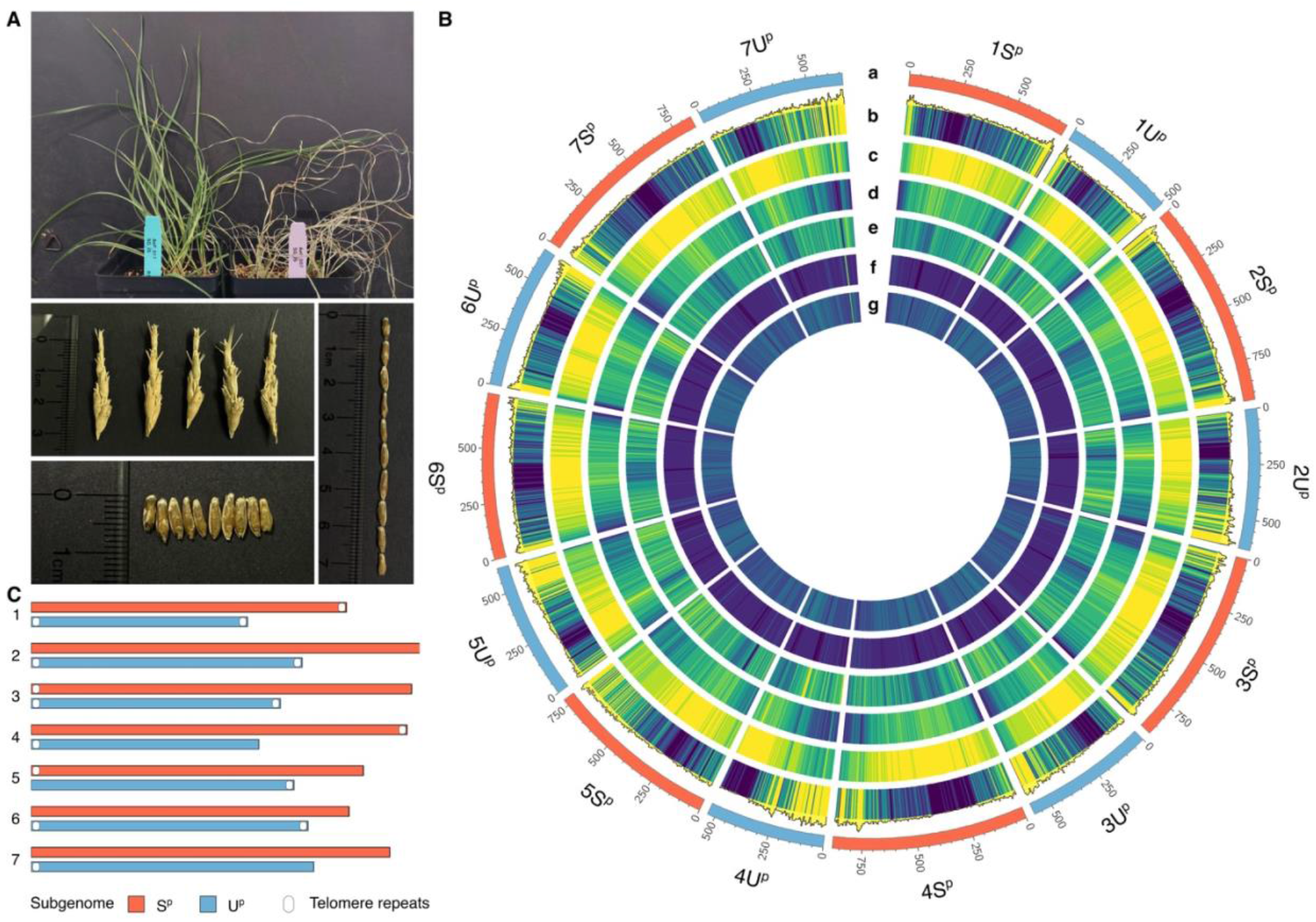
Phenotype and chromosome-scale reference genome assembly of *Aegilops peregrina*. **(A)** *Ae. peregrina* accession PI 604178 (left) showing seedling drought tolerant response in comparison to susceptible accession (right), together with representative spikes and seeds of PI 604178. **(B)** Circular representation of 14 chromosomes comprising seven S^p^ and seven U^p^ subgenomes. Tracks from outside to inside represent: (a) chromosomes, (b) gene density, (c) repeat density, (d) Gypsy retrotransposon diversity, (e) Copia retrotransposon density, (f) CACTA density, and (g) GC content distribution. **(C)** Telomeric repeat arrays detected at the chromosome ends of S^p^ and U^p^ subgenomes.

Here, we present a subgenome-resolved, chromosome-scale reference genome assembly of *Ae. peregrina* accession PI 604178 generated using PacBio HiFi and Hi-C sequencing data. The 10.13 Gb assembly contains 10.01 Gb (98.81%) anchored to 14 chromosome-scale pseudomolecules representing 1S^p^-7S^p^ and 1U^p^-7U^p^ chromosomes (Fig. 1B and 1C). The assembly has a contig N50 of 25.84 Mb and scaffold N50 of 746.48 Mb (Fig. 2A; Table 1). High quality of this assembly was supported by a consensus QV of 74.61, 97.83% k-mer completeness, and 99.9% complete BUSCOs. The independent BUSCO analyses of S^p^ and U^p^ subgenomes recovered 99.3% and 99.5% complete orthologs, respectively (Fig. 2B). LTR Assembly Index (LAI) values of 20.43 and 18.79 for S^p^ and U^p^, respectively, showed high continuity across repeat-rich regions in two subgenomes (Table 1). Repetitive sequences comprise 85.93% of the chromosome-anchored assembly, with Gyspy, Copia, and CATCA elements representing the predominant transposable element (TE) groups (Fig. 1B and 3A; Table 2). Structural gene annotations identified 59,910 high-confidence (HC) and 21,746 low-confidence (LC) protein coding genes (Fig. 3B; Table 3). We additionally assembled complete chloroplast and mitochondrial genomes in *Ae. peregrina*. This chromosome-scale reference genome establishes a genomic framework for the genetic and evolutionary study of *Ae. peregrina*. The resource will facilitate sequence-level characterization of disease and pest resistance loci introgressed into wheat, development of subgenome specific markers for tracking *Ae. peregrina* chromatin in wheat backgrounds, comparative analyses of S^p^ and U^p^ subgenomes with their diploid relatives, and investigation of genome evolution following allopolyploidization. It also expands the genomic resources available for exploiting *Aegilops* diversity in wheat improvement.

**Fig. 2:**
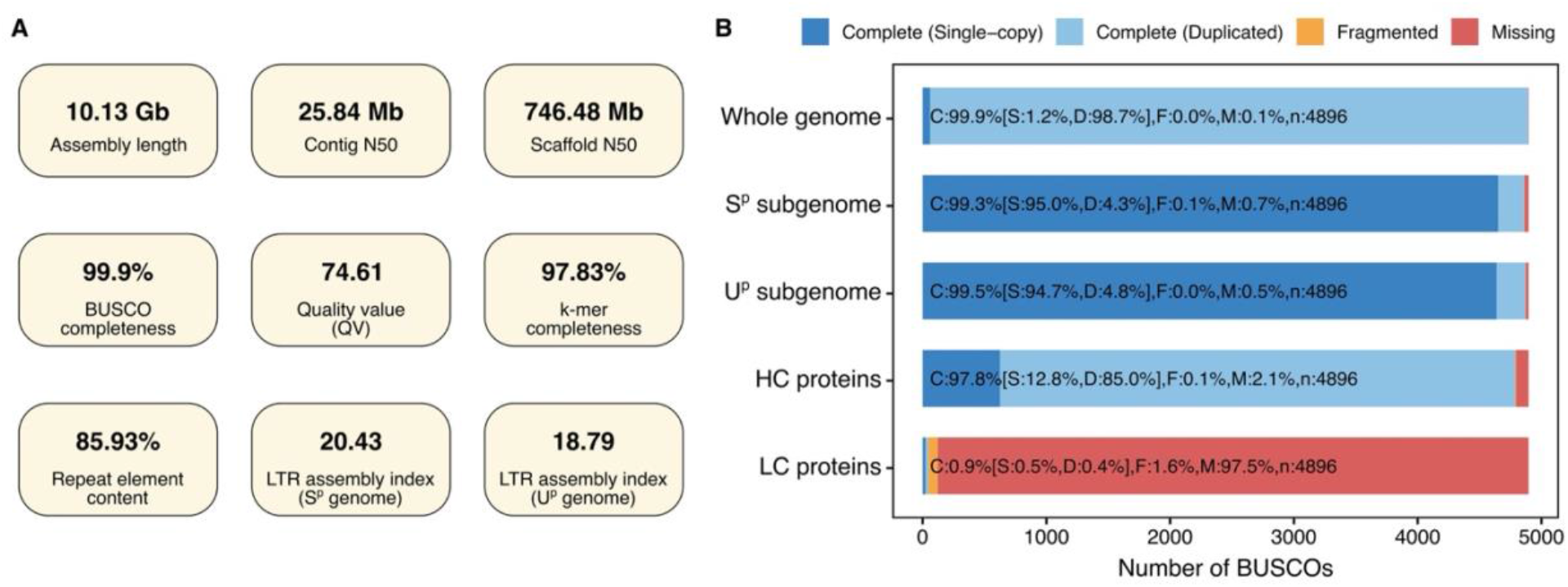
Assembly and annotation quality assessment of *Ae. peregrina* reference genome. **(A)** Summary of major assembly quality metrics, including total assembly length, contig N50, scaffold N50, BUSCO completeness, consensus quality value (QV), k-mer completeness, repeat content, and LTR Assembly Index (LAI) for the Sᵖ and Uᵖ subgenomes. **(B)** BUSCO assessment using the poales_odb10 dataset (n = 4,896) for the whole-genome assembly, independently evaluated Sᵖ and Uᵖ subgenomes, and high-confidence (HC) and low-confidence (LC) predicted protein sets.

**Fig. 3:**
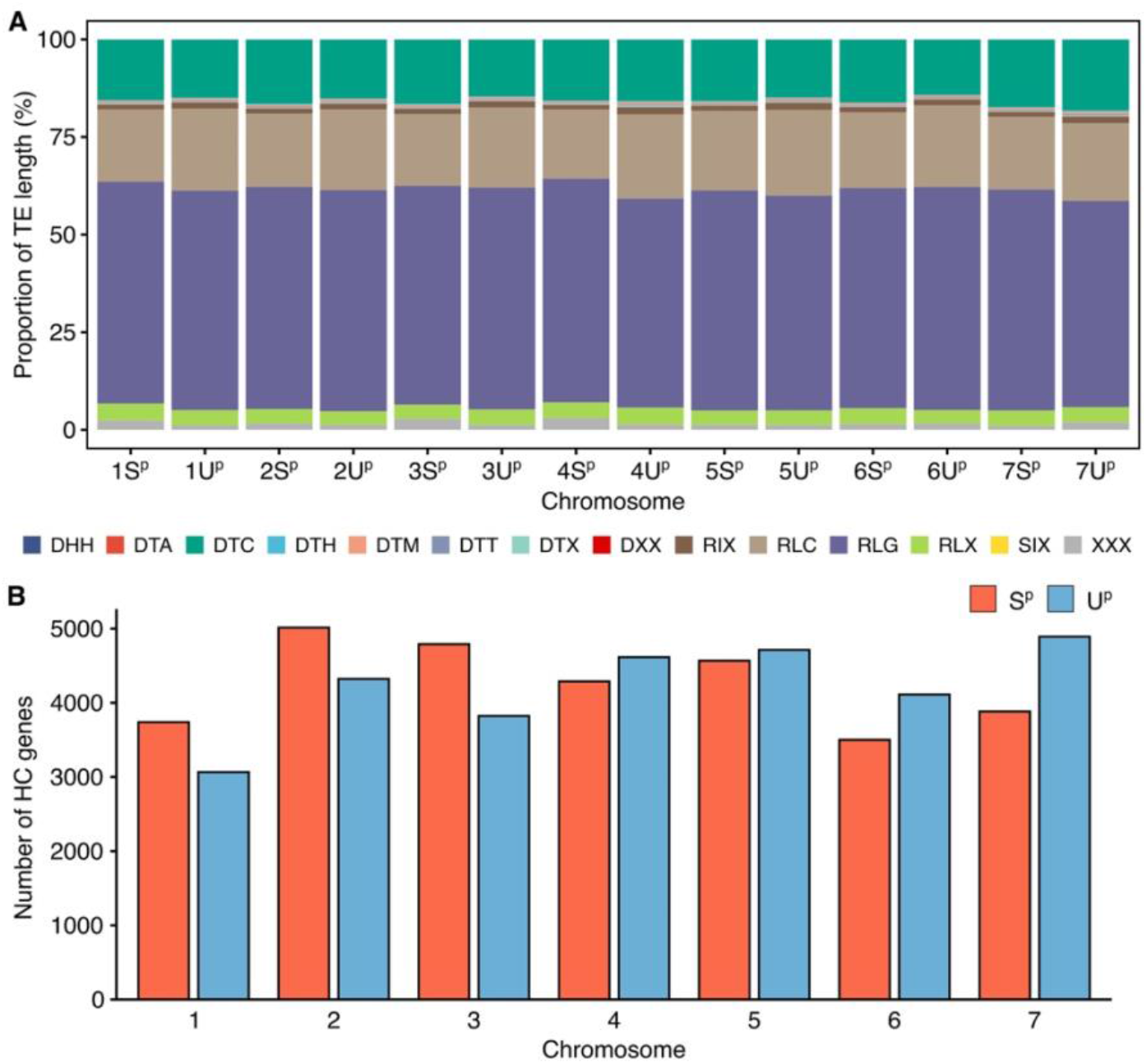
Transposable element composition and high-confidence gene distribution across the *Ae. peregrina* chromosomes. **(A)** Relative contribution of major transposable element classes to the repeat-annotated sequence of each Sᵖ and Uᵖ chromosome. **(B)** Number of high-confidence (HC) protein-coding genes on corresponding Sᵖ and Uᵖ chromosomes for each of the seven homoeologous groups.

**Table 1:**
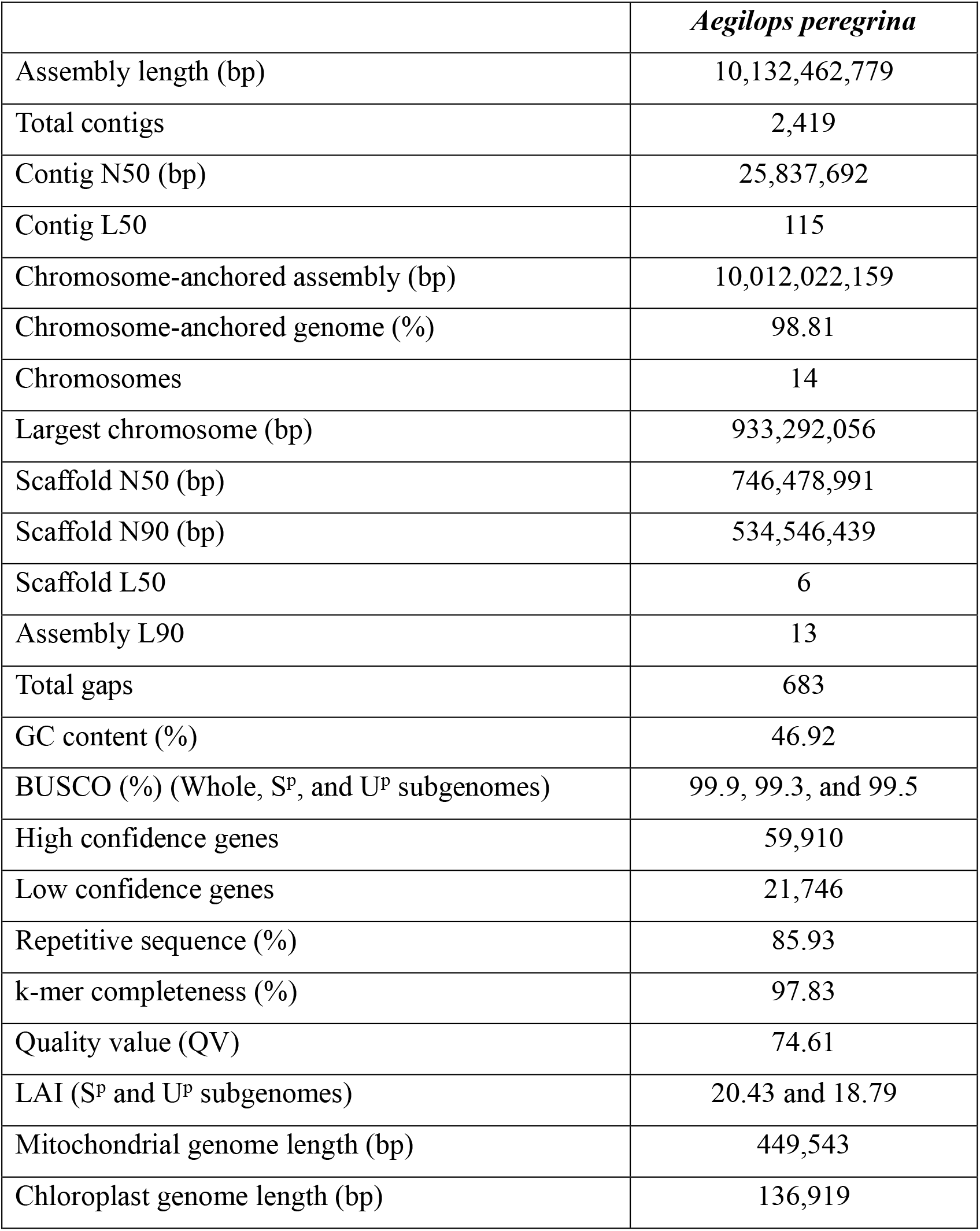
Assembly and quality statistics of *Aegilops peregrina* genome.

**Table 2:** Summary of transposable element (TE) composition in the *Aegilops peregrina* genome assembly.

|  | *Aegilops peregrina* |  |  |
| --- | --- | --- | --- |
|  | **Number of elements** | **Length (bp)** | **Genome proportion (%)** |
| All TEs | 3,211,306 | 8,603,114,080 | 85.93% |
| **Class 1** | 1,731,462 | 6,970,994,209 | 69.63% |
| *LTR retrotransposons* |  |  |  |
| Gypsy (RLG) | 1,083,766 | 4,832,078,711 | 48.26% |
| Copia (RLC) | 381,927 | 1,691,069,212 | 16.89% |
| Unclassified RT-LTRs (RLX) | 127,897 | 328,807,738 | 3.28% |
| *Non-LTR retrotransposons* |  |  |  |
| LINE (RIX) | 132,865 | 118,182,367 | 1.18% |
| SINE (SIX) | 5,007 | 856,181 | 0.01% |
| **Class 2** | 1,074,976 | 1,480,451,758 | 14.79% |
| *DNA transposons* |  |  |  |
| CACTA (DTC) | 688,408 | 1,363,594,975 | 13.62% |
| Mutator (DTM) | 92,486 | 45,947,755 | 0.46% |
| Harbinger (DTH) | 40,216 | 17,756,826 | 0.18% |
| Mariner (DTT) | 140,816 | 18,731,287 | 0.19% |
| Unclassified with TIRs (DTX) | 99,904 | 25,747,785 | 0.26% |
| Unclassified Class 2 (DXX) | 10,002 | 6,319,159 | 0.06% |
| hAT (DTA) | 1,530 | 1,066,261 | 0.01% |
| Helitrons (DHH) | 1,614 | 1,287,710 | 0.01% |
| **Unclassified repeats (XXX)** | 404,868 | 151,668,113 | 1.51% |

**Table 3:** Summary statistics of high confidence gene models in *Aegilops peregrina* genome.

| Features | S<sup>p</sup> subgenome | U<sup>p</sup> subgenome | Whole assembly |
| --- | --- | --- | --- |
| Total genes | 29,784 | 29,543 | 59,910 |
| Total transcripts | 32,851 | 32,406 | 65,912 |
| Total exons | 153,804 | 150,798 | 307,203 |
| Average gene length (bp) | 3,027 | 2,946 | 2,985 |
| Median gene length (bp) | 1,867 | 1,869 | 1,863 |
| Average CDS length (bp) | 1,295 | 1,292 | 1,293 |
| Median CDS length (bp) | 1,107 | 1,107 | 1,107 |
| Average exon length (bp) | 276.6 | 277.6 | 277.5 |
| Median exon length (bp) | 141 | 141 | 141 |
| Average intron length (bp) | 502 | 491.3 | 497.5 |
| Median intron length (bp) | 132 | 132 | 132 |
| Exons per gene | 5.16 | 5.1 | 5.13 |
| Monoexonic genes | 8,556 | 8,825 | 17,637 |
| Multiexonic genes | 21,228 | 20,718 | 42,273 |
| Exons per transcript | 4.68 | 4.65 | 4.66 |
| Monoexonic transcripts | 9,576 | 9,885 | 19,756 |
| Multiexonic transcripts | 23,275 | 22,521 | 46,156 |

Note: Whole assembly values also include gene models located on unplaced scaffolds.

## Methods

### Plant material and DNA extraction

We selected *Ae. peregrina* accession PI 604178 for whole genome assembly based on its seedling drought tolerance response (Fig. 1A) as per in-house developed protocol described in Gudi et al., 2026^21^ and resistance to stem rust races^20^, such as TTKSK and TRTTF. The seeds were procured from the United States Department of Agriculture-Germplasm Resources Information Network (USDA-GRIN). The accession PI 604178 was originally collected from Berekhya, Southern Coastal Plain, HaDarom, Israel (31.66666667, 34.63333333) with the sea elevation of 80 meters. The single seed decent method was used to increase seeds for this accession and to maintain genetic purity. Plants were grown in greenhouse under 16 hours (h) of daylight with 22°C day and 18°C night temperature. Young leaves collected from three-week old plants were immediately frozen in liquid nitrogen and stored at-80°C until DNA extraction. The high molecular weight (HMW) DNA was extracted using a CTAB-Qiagen Genomic-tip method^22^. We used nanodrop spectrophotometer and Qubit dsDNA HS assay to assess the quality and quantity of the extracted HMW DNA before library preparation.

### PacBio HiFi sequencing and genome assembly

The HMW DNA was sheared to 17 kb fragments with Diagenode Megaruptor and SMRTbell Express Template Prep kit 2.0 was used for PacBio libraries^23^. The fragments larger than 10 kb were selected using Sage BluePippin size selection kit and were sequenced on the PacBio Revio platform using three 30 h SMRT cells. The Sequel II Sequencing kit 2.0, Sequencing Primer v5, and Sequel Binding kit 2.2 were used. The HiFi sequencing generated 217.62 Gb of data with an average read length of 14,782 bp and a median read quality score of Q33 (∼99.95% accuracy). Approximately 21.5x PacBio HiFi genome coverage was produced. The genome size and k-mer characteristics were estimated using GenomeScope2 (v2.0.1)^24^. The k-mer frequency distributions were generated from HiFi data using Meryl (v1.4.2)^25^. GenomeScope2 analyses were performed at k-mer lengths of 31 and 41. At k=31, GenomeScope2 estimated a genome size of approximately 9.97 Gb with unique sequence fraction of 23.4%, and a k-mer error rate of 0.0912%. Analysis at k=41 produced a genome size estimate of approximately 10.01 Gb, with 31.6% unique sequence and a k-mer error rate of 0.105% (Fig. 4A). Estimated heterozygosity was approximately 0.001% in both analyses. The k-mer spectrum contained a major peak at approximately 21x coverage.

**Fig. 4:**
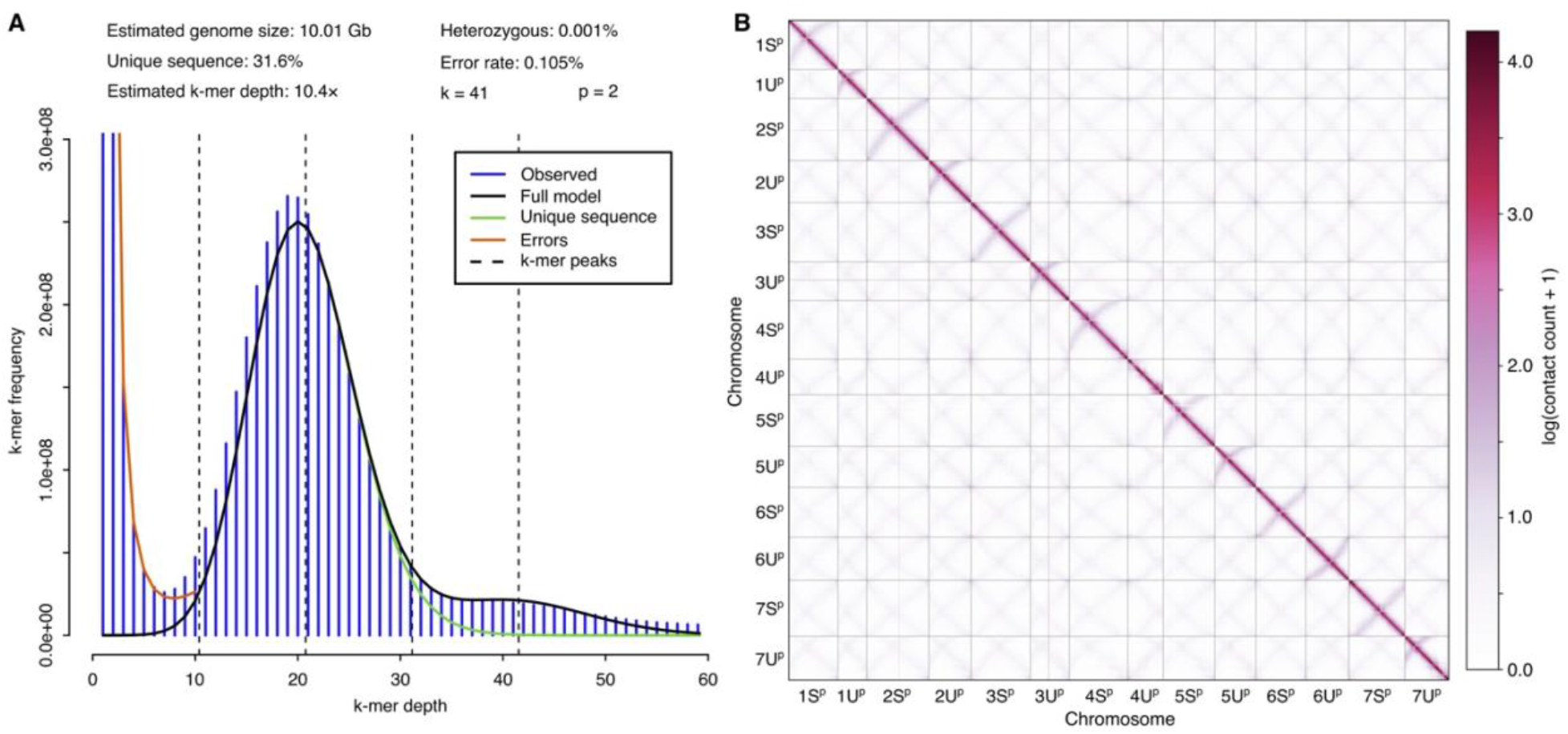
Genome k-mer characteristics and Hi-C validation of chromosome-scale *Ae. peregrina* assembly. **(A)** GenomeScope2 k-mer frequency distribution generated from PacBio HiFi reads at k = 41. **(B)** Hi-C contact matrix of 14 chromosomes at 250 kb resolution.

We used hifiasm (v0.25.0-r726)^26^ to assemble PacBio HiFi reads into contigs with default parameters for an inbred species. The primary assembly consisted of 2,419 contigs spanning 10.19 Gb, with an average contig length of 4.22 Mb, and a largest contig length of 111.49 Mb. The assembly had a contig N50 of 25.84 Mb with a contig L50 of 115, indicating high continuity prior to Hi-C scaffolding (Table 1). The primary contig assembly contained no gaps prior to scaffolding.

### Organelle de-novo assembly

Chloroplast and mitochondrial genomes were assembled independently from PacBio HiFi reads using Oatk (v1.0)^27^. The parameters “-k 1001-c 100-m embryophyta_mito.fam-p embryophyta_pltd.fam” were used. The chloroplast and mitochondrial genomes were assembled as single circular genome, with the size of 136,919 bp and 449,543 bp, respectively.

### Hi-C sequencing and genome scaffolding

We generated 1,206,141,041 chromosome conformation capture (Hi-C) read pairs for scaffolding of primary contig assembly. The leaf tissue from the same three-week old plant used for PacBio sequencing were collected and submitted to Phase Genomics (Settle, WA) for Hi-C library preparation. Three Hi-C libraries were prepared following the standard Phase Genomics proximity ligation protocol. The crosslinked chromatin was digested with four-enzyme cocktail (DpnII, DdeI, HinfI, and MseI), subjected to proximity ligation, and prepared as 2 x 150 bp paired-end sequencing libraries for Illumina sequencing. The raw Hi-C reads were aligned to the primary contig assembly following the Arima Mapping Pipeline (A160156_v03, Arima Genomics, Carlsbad, California, USA). The mapping showed a high alignment rate with 99.06% of read pairs having both mates mapped to primary assembly. Duplicate alignments were identified using SAMBLASTER (v0.1.26)^28^, which marked 657,857,611 of the mapped pairs as duplicates. After filtering and duplicate removal, 536,891,087 non-duplicate Hi-C read pairs were retained for scaffolding. We used a proximity guided scaffolder, YaHS (v1.2a.2)^29^, for scaffolding of our primary assembly and found 28,928,874 intra-chromosomal and 507,962,213 inter-chromosomal contacts. The manual inspection and correction was performed using JuiceBox (v2.15)^30^. The inter-contig gaps within the scaffolds were padded with 100Ns. A final Hi-C contact matrix was generated at 250 kb resolution.

### Assembly quality assessment

To screen the *Ae. peregrina* assembly for any contaminants, FCS-GX^31^ with “fcs.py screen genome” and “fcs-gx” database with *Ae. peregrina* taxonomic identifier 130461 was used, while adapter screening was performed with “run_fcsadaptor.sh --euk”. No contaminants or adapter sequences were reported in the screening check. The de-novo assembled organelles were mapped to the scaffolded assembly using minimap2 (v2.30-r1287)^32^ to identify mitochondrial and chloroplast sequences. The analysis identified 1,408 organelle-derived scaffolds, which were filtered out from the assembly. Additionally, short contigs (<20 kb) were excluded from the final assembly. In total, 1,478 scaffolds totaling 66.45 Mb were removed from the final nuclear genome assembly. The final assembly contained 614 scaffolds spanning 10.13 Gb, of which 10.01 Gb (98.81%) was assigned to 14 chromosome-level pseudomolecules representing seven S^p^ and seven U^p^ chromosomes (Fig. 1B and Table 1). The remaining 120.44 Mb was retained as unplaced scaffolds. For naming and orienting the pseudomolecules, *Ae. longissima* accession AEG-6782-2 (v2)^33^ (S genome) and *Ae. umbellulata* accession PI 554389^34^ (U genome) assemblies were used. We performed a telomere search on all 14 chromosomes using tidk (v0.2.31)^35^ with “clade” option set to “Poales”. The chromosome-level genome assembly of *Ae. peregrina* PI 604178 was evaluated for accuracy, completeness, and contiguity, using tools such as QUAST (v5.2.0)^36^, assembly-stats (v1.0.1), Merqury (v1.4.1)^25^, BUSCO (v 6.0.0)^37^, and LTR_retriever (v3.0.4)^38^. The pairwise dotplots were generated with chromeister (v1.5.a)^39^. The plots in this study are generated with ggplot2^40^ package in R (4.4.0) and figure panels were arranged in Inkscape.

### Structural and functional annotations

We used well curated ClariTeRep database^41^ for transposable element (TE) annotations. The RepeatMasker (v4.1.5)^42^ was run using “-xsmall-xm-engine rmblast” parameters and the output was processed with “clariTE.pl” script (https://github.com/jdaron/clari-te) to merge fragmented matches, reconstruct nested insertions, resolve element boundaries and assign TE copies according to the ClariTeRep hierarchical classification. TE abundance and sequence coverage were calculated for 14 chromosomes (Table 2).

The soft-masked genome was used for structural annotations of gene models using BRAKER3^43^. We used BRAKER3 in ETP mode and integrated transcript, protein-homology, and *ab initio* evidence for protein-coding gene annotations. The transcript evidence consisted of publicly available *Ae. peregrina* RNA-seq datasets SRR16550937-SRR16550944^citation^. The “viridiplantae” (https://bioinf.uni-greifswald.de/bioinf/partitioned_odb12/Viridiplantae.fa.gz) protein evidence were downloaded from OrthoDB (v12)^37^. The predicted protein coding genes were functionally annotated with InterProScan (v5.38-76.0)^44^ and putative gene functions were assigned in the gff3 files, based on the domain conserveness and sequence similarity with curated Unisprot-Sprot protein database (https://www.uniprot.org/help/downloads). The gene identifiers follow the format “AeP.PI604178.r1.[chromosome]G[GeneID]. For instance, “AeP.PI604178.r1.1SpG0000100” denotes a gene located on chromosome 1S^p^ of the PI 604178 release 1 annotation. We categorized the gene models into high-confidence (HC) and low-confidence (LC) groups based on the sequence similarity with the Magnoliopsida UniProt TrEMBL dataset and the Triticeae repetitive element database (TREP). HC genes were defined as those sharing significant similarity (E-value < 1e-10) with TrEMBL proteins, demonstrating > 66% amino acid identity and < 25% difference in sequence length^45^. Those gene models that missed these homology thresholds, lacked external support, or matched TE sequences in TREP database (E-value < 1e-10), were classified as LC. The chromosome-anchored HC annotation comprised 29,784 genes on the S^p^ subgenome and 29,543 genes on the Uᵖ subgenome. The gene architecture was highly similar between the two subgenomes (Table 3).

### Data Records

The genome assembly for *Ae. peregrina* accession PI 604178 is deployed to GrainGenes Genome Browser (https://graingenes.org/GG3/genome_browser)^46^ with BLAST functionality. We deposited raw PacBio HiFi sequencing data generated for *Ae. peregrina* accession PI 604178 in NCBI Sequence Read Archive (SRA) with accession numbers SRR40310784, SRR40310785, and SRR40310786. The Hi-C sequencing data was submitted to SRA with SRR40310783 accession number. Genome has been deposited at DDBJ/ENA/GenBank under the accession JCCJTD000000000. The version described in this paper is version JCCJTD010000000. The reference genome assembly is accessible through GenBank under GCA_########.1as a part of BioProject PRJNA1517304. Datasets such as mitochondrial and chloroplast genomes, soft-masked genome assembly, repeat annotations, structural gene models, and functional gene annotations are available via figshare.

### Technical validation

The Hi-C contact map showed strong and continuous intra-chromosomal interaction signals across all 14 pseudomolecule diagonals, supporting accurate chromosome-scale contiguity (Fig. 4B). FCS-GX screening detected no foreign sequence contamination or adapter sequences. Merqury analysis showed a consensus QV of 74.61 and k-mer completeness of 97.83% (Fig. 2A). The k-mer copy-number spectrum showed that most reliable read-derived k-mers were represented in the assembly, with the predominant assembly-specific peak corresponding to single-copy representation (Fig. 5A). Chromosome-specific QV ranged from 77 to 82, demonstrating consistently high consensus accuracy across both S^p^ and U^p^ chromosomes (Fig. 5B). Telomeric repeat arrays were identified at 4 of 14 chromosome ends in the Sᵖ subgenome and 10 of 14 chromosome ends in the Uᵖ subgenome (Fig. 1C). Finaly, LAI values of 20.43 for Sᵖ and 18.79 for Uᵖ further supported high continuity of repeat-rich regions. Whole genome pairwise comparisons showed strong chromosome-level collinearity of the Sᵖ subgenome with *Ae. longissima* and the Uᵖ subgenome with *Ae. umbellulata*, supporting the assignment, ordering, and orientation of the 14 *Ae. peregrina* chromosomes (Fig. 6). The quality of the structural gene annotation was independently evaluated using BUSCO analysis of the HC protein set, which recovered 97.8% complete BUSCOs, with 0.1% fragmented and 2.1% missing orthologs, supporting the high completeness of the predicted protein-coding gene space (Fig. 2B). In contrast, the LC protein set recovered only 0.9% complete BUSCOs, consistent with the enrichment of conserved protein-coding gene space within the HC annotation. The HC gene annotation contained 59,910 genes, distributed nearly equally between the S^p^ (29,784) and Uᵖ (29,543) subgenomes. Gene structural features were also highly comparable between the two subgenomes, including gene and CDS lengths, exon and intron lengths, and exon number per gene (Table 3), indicating consistent annotation quality across both subgenomes.

**Fig. 5:**
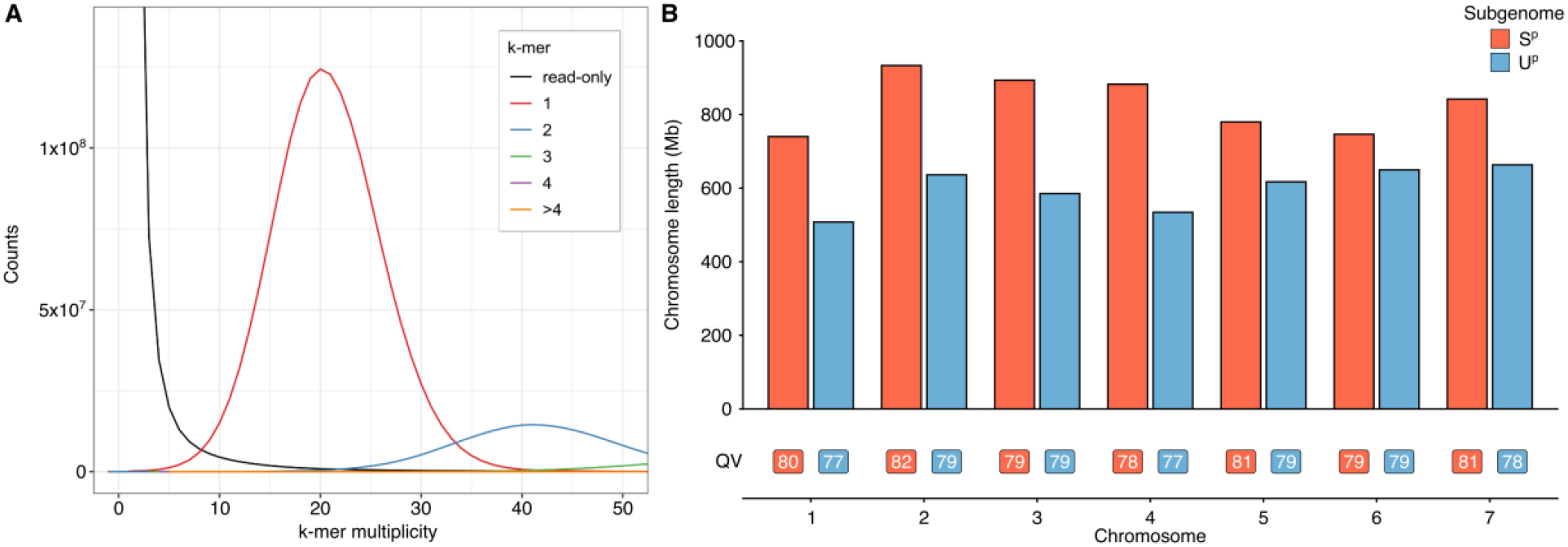
k-mer based evaluation of assembly completeness and chromosome-level consensus accuracy. **(A)** Merqury k-mer copy-number spectrum comparing PacBio HiFi read-derived k-mers with their representation in the final assembly. Curves represent read-only k-mers and k-mers occurring one, two, three, four, or more than four times in the assembly. **(B)** Lengths and Merqury consensus quality values (QV) of corresponding Sᵖ and Uᵖ chromosomes across the seven homoeologous groups.

**Fig. 6:**
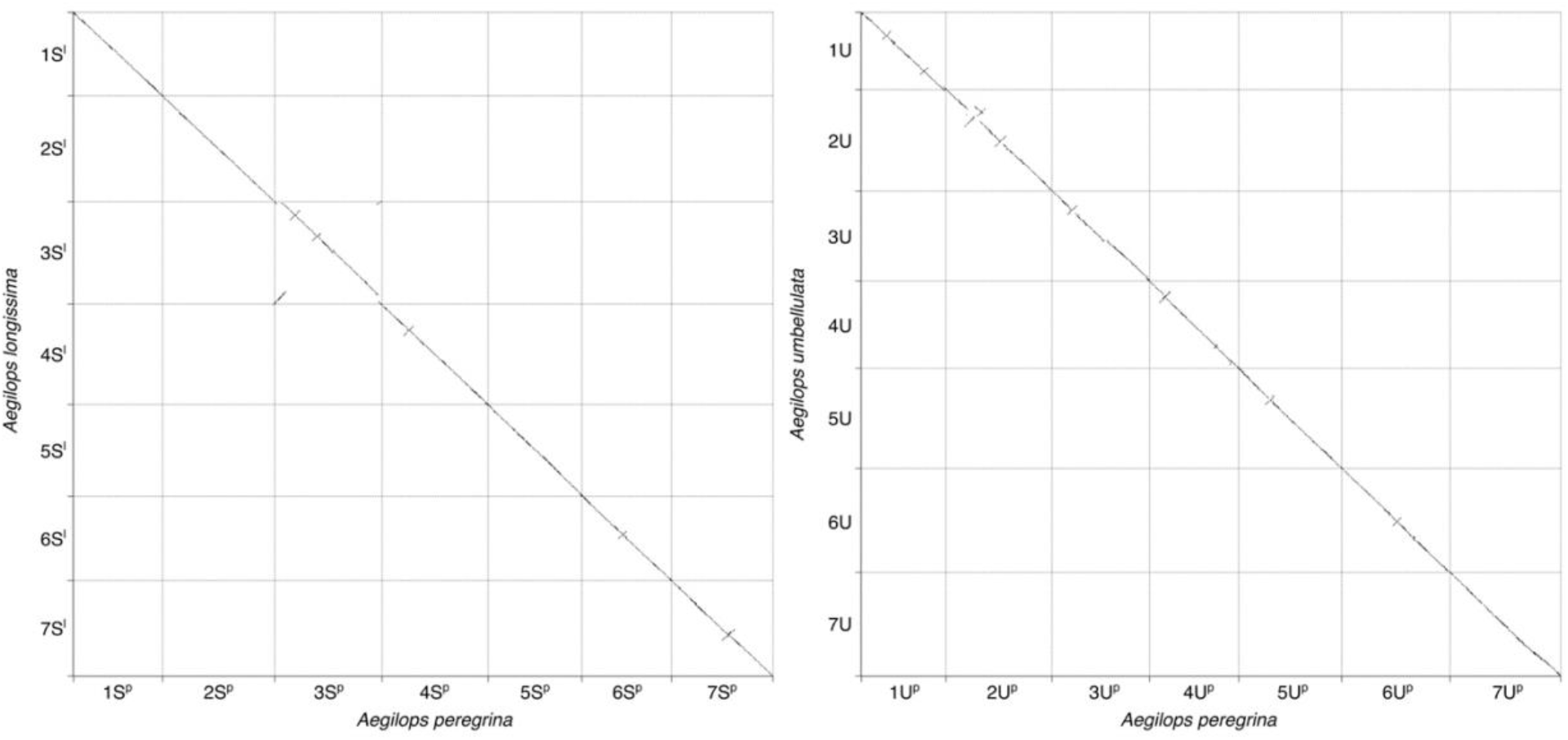
Chromosome-level collinearity of *Ae. peregrina* Sᵖ and Uᵖ subgenomes with diploid relatives. Whole genome dotplot comparisons of the seven *Ae. peregrina* Sᵖ chromosomes with the S genome chromosomes of *Ae. longissima* (left) and the seven Uᵖ chromosomes with the U-genome chromosomes of *Ae. umbellulata* (right). Continuous chromosome-scale diagonals demonstrate extensive macro-collinearity and support the assignment, ordering, and orientation of the *Ae. peregrina* pseudomolecules.

### Usage Notes

To allow community access, the *Ae. peregrina* genome assembly and annotations have been submitted to public repositories. Interactive genome viewer for this assembly is available through GrainGenes genome browser. The direct BLAST functionality is also available at https://graingenes.org/blast/.

## Data Availability

The raw sequencing data, genome assembly, and gene model annotations have been deposited in public repositories. The data generated for *Aegilops peregrina* accession PI 604178 is submitted to NCBI under BioProject accession number PRJNA1517304 (https://www.ncbi.nlm.nih.gov/bioproject/PRJNA1517304/). The raw Hi-C and PacBio HiFi sequencing data is accessible through SRA accession numbers from SRR40310783 to SRR40310786 (https://identifiers.org/ncbi/insdc.sra:SRP729778). The genome assembly is made available on GenBank, figshare, and GrainGenes Browser. The supporting datasets including gene annotations and organelle genome assemblies are accessible via figshare (https://doi.org/10.6084/m9.figshare.33330000).

## Code Availability

In this study, we used publicly available bioinformatics tools, and no custom scripts were developed. We used default software parameters, and detailed information on software names and versions is provided in the Methods section. All command-line procedures and analyses can be reproduced using the listed software and parameters.

## Author Contributions

R.G. and J.S conceived and designed the study. J.S., S.G., and P.J.M. performed data analyses. J.S. prepared the first draft of the manuscript. J.S., S.G., P.J.M, U.G., and R.G. contributed to data generation, interpretation of results, and manuscript editing. R.G. procured the funding and supervised the project. All authors reviewed and approved the final manuscript.

